# In search of the universal method: a comparative survey of bottom-up proteomics sample preparation methods

**DOI:** 10.1101/2022.05.04.490597

**Authors:** Gina Varnavides, Moritz Madern, Dorothea Anrather, Natascha Hartl, Wolfgang Reiter, Markus Hartl

## Abstract

Robust, efficient and reproducible protein extraction and sample processing is a key step for bottom-up proteomics analyses. While many sample preparation protocols for mass spectrometry have been described, selecting an appropriate method remains challenging, since some protein classes may require specialized solubilization, precipitation, and digestion procedures. Here we present a comprehensive comparison of 16 most widely used sample preparation methods, covering in-solution digests, device-based methods, as well as commercially available kits. We find a remarkably good performance of the majority of the protocols with high reproducibility, little method dependencies and low levels of artifact formation. However, we revealed method-dependent differences in the recovery of specific protein features, which we summarized in a descriptive guide-matrix. Our work thereby provides a solid basis for the selection of MS sample preparation strategies for a given proteomics project.

## Introduction

State-of-the-art mass-spectrometry-based proteomics workflows are sophisticated multi-step processes, combining different methodologies and instrumentation. Clearly, the data quality of an experiment depends on the characteristics and limitations of each step, with errors or biases propagating from the first step throughout the whole experiment. For this reason, the sample preparation protocol is a key determinant in defining what proportion of the proteome is available for the ensuing analysis. Moreover, the robustness and reproducibility of this step will define the degree of data variation and potential systematic bias. Ideally, the universal sample preparation protocol would efficiently and robustly isolate all proteins of any given sample to near completeness. In reality, such a comprehensive isolation is very challenging as proteins constitute a heterogeneous group of macromolecules in terms of physicochemical properties, and subcellular localization. In addition, other sample characteristics, such as rigid cell walls and tissues that are difficult to lyse or interfering cellular components (e.g. nucleic acids, metabolites, etc.), can greatly affect isolation efficacy and analysis and need to be addressed.

To solve these problems, different sample preparation methods have been developed that can be divided into in-solution digestion methods and methods using additional devices such as filters or beads for protein immobilization or purification, or both. Classical in-solution digestion (ISD) protocols essentially differ in the choice of buffer systems, which are either based on chaotropic denaturants, such as urea or guanidine hydrochloride (GnHCl), or surfactants, such as the ionic detergent sodium-dodecyl-sulfate (SDS) or the bile salt sodium-deoxycholate (SDC), as they effectively solubilize and denature proteins ^1–3^. Recently, a novel ISD strategy, Sample Preparation by Easy Extraction and Digestion (SPEED), has been published ^4^which uses neither detergents nor chaotropic agents for protein extraction but which is solely based on dissolving proteins in trifluoroacetic acid (TFA). Further, ISD protocols often require protein precipitation using either acetone ^5^, alcohols such as ethanol ^6^, or chloroform/methanol ^7^, in order to avoid carry-over of non-protein components that might interfere with downstream processing or analysis.

Device-based approaches (hereafter referred to as “cleanup methods”) aim to remove interfering substances before digestion in ‘‘reactors” or on beads. For example, Filter-Aided Sample Preparation (FASP) ^8^, utilizing molecular weight cut-off (MWCO) membranes, and suspension trapping (S-Trap) ^9^, applying three-dimensional porous quartz filter materials, capture proteins on filters enabling detergent removal, protein digestion, and peptide recovery. Single-pot, solid-phase-enhanced sample preparation (SP3) ^10^ (and also SP4 ^11^), uses on-bead based purification and digestion of proteins in a single tube, exploiting the property of denatured proteins to be non-specifically immobilized on microparticles by protein aggregation ^12^. Finally, the original in-StageTip (iST) method utilizes C18 discs prepared in pipette tips or cartridges, to trap proteins for digestion and subsequently to desalt the peptides ^13^. Based on these and similar methodical concepts, commercially available MS sample preparation kits in different formats have been developed for iST (PreOmics), S-Trap (ProtiFi), and in-solution digests coupled to peptide cleanup columns (EasyPep, Thermo Scientific).

Overall, this almost overwhelming number of protocols and variants with their apparent advantages and disadvantages make the selection of a suitable method for a given project difficult. Although previous studies compared selected sets of protocols, often focusing on particular aspects or on presenting a new method, a comparison including the most commonly used in-solution, device-based, and commercial methods had yet to be conducted ^1,3,4,12,14–18^. It is also debatable, whether there is a truly universal method that exhibits no or negligible extraction bias, as has been proposed for some protocols ^4,17^, and which is applicable to all types of samples. Proving universality is an almost futile task, as it would require the comparison of a set of methods for a virtually endless list of cell types, tissues, body fluids, and organisms. However, Glatter *et al.*^1^ and Doellinger *et al.*^4^ convincingly demonstrated for a selection of ISD protocols and device-based protocols that there are organism- and buffer-specific differences in extraction efficiency, when comparing samples of different bacterial and human cell line. From these studies, it can be expected that such differences will further increase when comparing even more diverse sets of sample types, e.g. mammalian tissues, plants, or fungi. In contrast, investigating differences in proteome composition for a given set of methods in a defined sample type is more feasible and allows to answer whether and how protocols differ in their extraction properties for the given sample type. In combination with more practical considerations, like for example processing time, ease of use, and consumable costs, this could help in making a more informed decision for a particular sample preparation strategy and serve as a blueprint for similar studies in other sample types.

Here, we prepared proteomes from HeLa cells applying classical ISD protocols based on urea-, GnHCl- and SDC-based buffer systems as well as SPEED ^4^, FASP ^8^, S-Trap ^9,19^ (ProtiFi), SP3 ^10^ protocols and two commercial kits; iST ^13^ (PreOmics) and EasyPep (Thermo Scientific). We therefore present a comprehensive quantitative and qualitative comparison of 16 of the most widely used MS sample preparation methods. Our experimental design maximizes reproducibility, comparability, and allows for unbiased statistical analyses to extract differences between the methods. The individual methods show a similar proteome extraction efficacy and coverage, based on identified proteins and peptides. Method-induced peptide artifacts seem to be negligible. However, an exploratory analysis based on k-means clustering revealed qualitative differences in extracted proteomes, which we mapped to features derived from protein databases. The results were summarized into a descriptive guide-matrix which highlights specific enrichment of protein features such as structure, abundance, and localization for individual methods. Consequently, our study provides a solid comparison of the currently most widely used sample preparation in proteomics and can be used as an aid in selecting MS sample preparation strategies.

## Results & Discussion

### Experimental design and quality control

To provide a comparative analysis of MS sample preparation methods, we applied 16 widely used protein extraction protocols to isolate whole-cell proteomes from HeLa cells and compared their efficacy on the basis of quantitative and qualitative parameters (**Figure 1A**). Our experimental setup covered in-solution digest (ISD) protocols as well as cleanup methods, including commercially available MS sample preparation kits.

**Figure 1.**
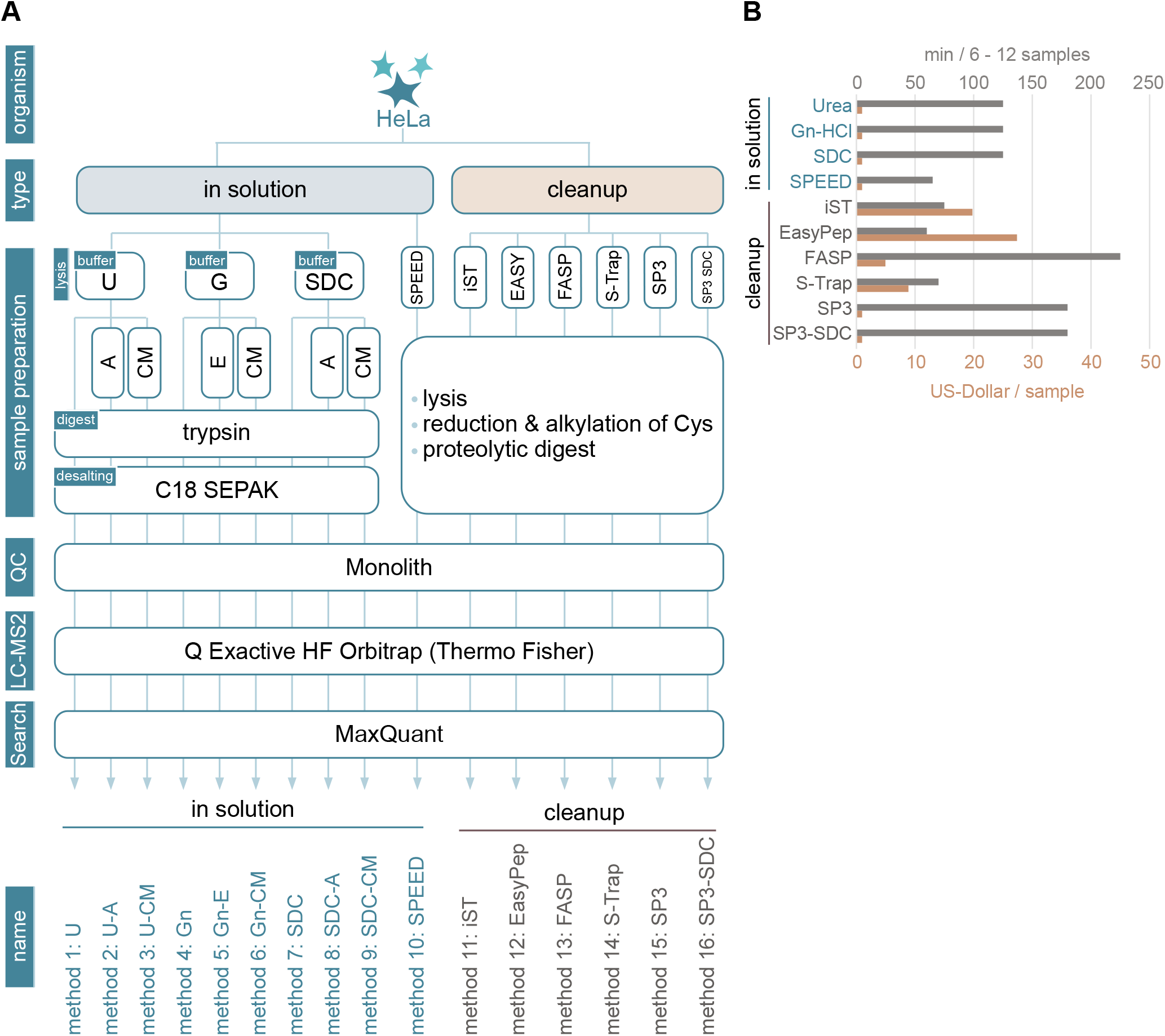
Experimental Setup and Quality Control of Applied MS Sample Preparation Methods. **(A)** Scheme of experimental design. Proteomes of HeLa cells were prepared according to indicated sample preparation methods. In solution digest (ISD) protocols (left) covered classical approaches based on urea (U)-, guanidine hydrochloride (G)-, or sodium deoxycholate (SDC)-buffered systems as well as Sample Preparation by Easy Extraction and Digestion ^4^ (SPEED). Lysates prepared with classical ISD protocols were either directly submitted to tryptic digestion or previously mixed with appropriate amounts of acetone (A), ethanol (E) or chloroform/methanol (CM) to precipitate proteins. Cleanup methods covered device-based approaches such as FASP, S-Trap, iST (PreOmics), EasyPep (Thermo Scientific), as well as SP3-based methods (right). Quality control (QC): peptide concentrations of all samples were determined using UV chromatograms (Monolith) after proteolytic digestion and adjusted for a concentration of 100ng/μl before MS analysis. All samples were analyzed using a quadrupole-orbitrap hybrid MS instrument MS raw data was analyzed using MaxQuant. **(B)** Overview of the average cost in US-Dollars (USD) per sample (brown) and time in minutes needed to process 6-12 samples (gray) excluding digestion and optional C18 peptide cleanups.

For classical ISDs, cells were lysed either at room temperature (urea) or 60°C (GnHCl, SDC) to optimize cell lysis and protein solubilization. We observed similar protein extraction efficiencies for the three applied buffer systems (**Supplemental Figure 1A**). Lysates were split into three groups of aliquots of equal protein amounts. The first group was directly subjected to proteolytic digestion using trypsin. The protein fractions of the remaining aliquots were additionally purified prior to proteolysis by acetone-, ethanol-(given that GnHCl is not soluble in acetone), or chloroform/methanol protein precipitation, respectively (**Figure 1A**). Notably, some combinations of buffer systems and precipitation methods, such as urea-based buffer and chloroform/methanol precipitation, resulted in significant sample losses. Highest yields were observed with SDC-based buffers (**Supplemental Figure 1A**) which corresponds to previous observations ^18,20^.

Cleanup samples were prepared as previously described ^8–10,13,19^ or according to manufacturer’s guidelines, with the exception of SP3, where additionally to the detergent-heavy buffer system, an easy to prepare buffer consisting of 1% SDC in Tris-HCl (see Methods for further information) was tested (SP3-SDC). The latter was included since this buffer composition delivered high performance in classical ISDs ^1,18^. Overall, we obtained similar peptide concentrations after tryptic digestion in all cleanup samples (**Supplemental Figure 1B**), even though proteolysis differed in reaction mix composition, reaction time, and peptide to enzyme ratio (see *Materials* and *Methods*).

To achieve equal loading for MS measurements, peptide concentrations of all samples were determined using UV chromatogram peak areas and adjusted accordingly. MS measurements were performed on a quadrupole-orbitrap hybrid MS instrument (**Figure 1A**). All 16 experimental conditions were analyzed in three technical replicates, resulting in a total of 48 MS runs which were measured in six consecutive batches. The performance of the LC-MS system was monitored by inspecting retention-times, intensities and peak shapes of spike-in standards (iRT) to ensure similar conditions within and between batches. Non-normalized summed protein group intensities indicated that comparable amounts of peptides were submitted to MS measurement (**Supplemental Figure 1C**).

### Cost and time effort

Since the expenditure of time and money is important to consider, we determined the average cost in US Dollars and hands-on sample processing times for the applied methods (**Figure 1B**). ISD protocols come at very low consumable costs but are, with the exception of SPEED, considerably more time demanding than commercial kits. EasyPep, iST, SPEED and S-Trap protocols were found to have similar hand-on times of around 60 min. FASP, on the other hand, is inherently more time-consuming with long centrifugation steps, taking up to four hours. Costs ranged from 1$ (ISD, SPEED, SP3 and SP3-SDC) to 5$ (FASP), ~10$ (S-Trap), ~20$ (iST) or ~30$ (EasyPep) per sample. From this perspective, SPEED represents a competitive protocol that combines short handling times with low consumable costs.

### Global comparison of performance

We first compared overall method performance, considering total numbers of protein groups (protein IDs) and peptides (peptide IDs) identified by LC-MS/MS (**Figure 2A, Supplemental Table 1**). After filtering data (see Materials & Methods), we retrieved protein IDs ranging from 3500 to 4500 and peptide IDs ranging from 30000 to 40000, with SDC-based sample preparations resulting in the highest numbers. ISD protocols based on GnHCl, on the other hand, delivered the lowest numbers of identified peptides and proteins.

**Figure 2.**
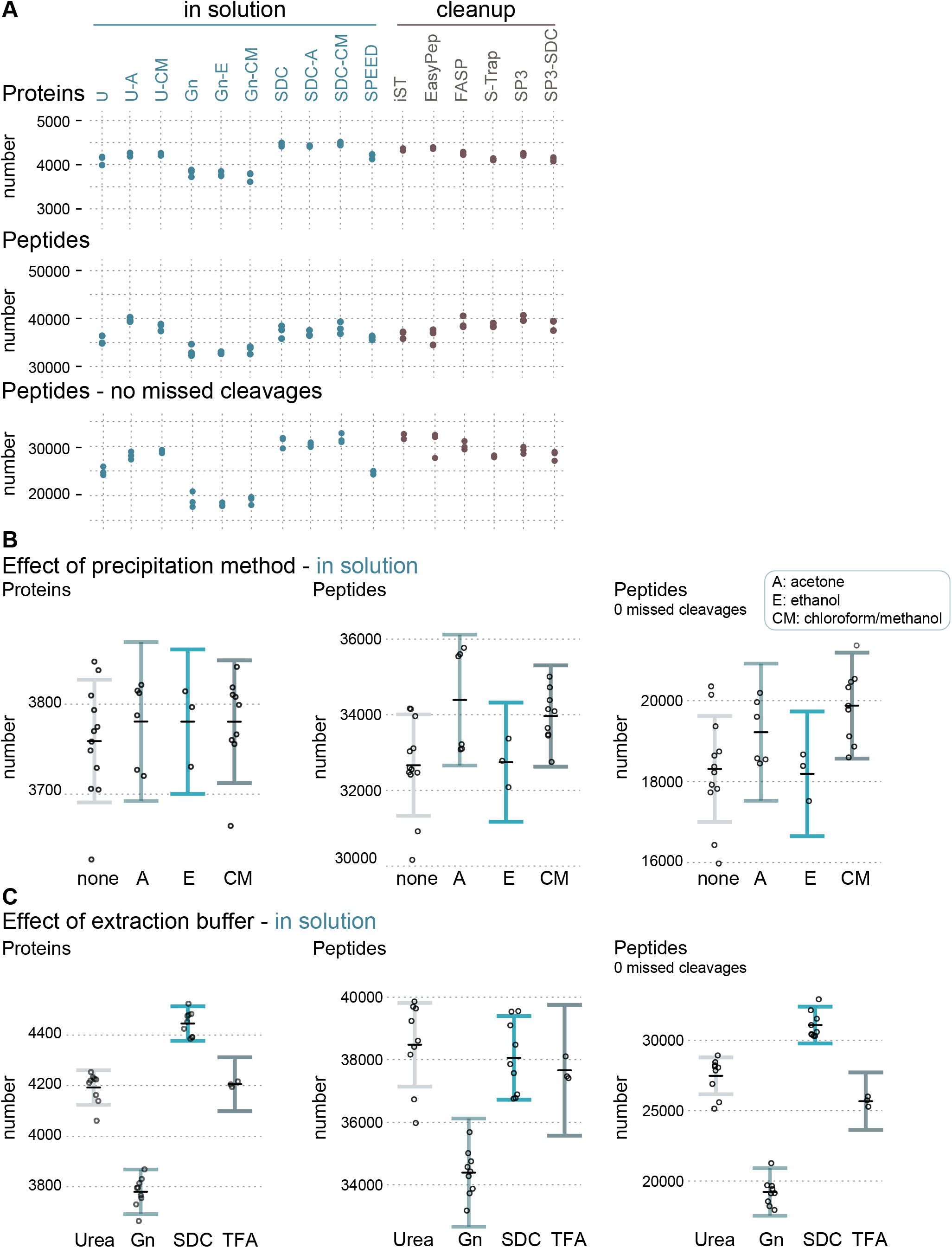
Partial Residual Plots highlighting effects of applied buffer systems and protein precipitation methods in ISD protocols. **(A)** Diagram showing a comparison of total number of identified (by MS/MS) proteins (top), peptides (middle) and peptides with no missed cleavages (bottom). **(B)** Effects of applied protein precipitation method on the number of identified (by MS/MS) proteins (left), peptides (middle) and peptides with no missed cleavages (right). A: acetone precipitation, E: ethanol precipitation, CM: chloroform-methanol precipitation. **(C)** Same as (B) except the effects of applied buffer systems is shown. Gn: guanidine hydrochloride, SDC: sodium-deoxycholate, TFA: SPEED ^4^. Data points represent the predicted number of IDs. Error bars correspond to a 95% confidence interval (CI). Black lines indicate the average predicted number of IDs.

Extraction buffers containing chaotropes or detergents are known to interfere with the protease activity of trypsin ^21,22^, which results in incomplete protein digestion and consequently in lower proteome coverage due to oversampling of different cleavage forms of abundant peptides. An analysis of missed cleavage frequencies clearly demonstrates strong differences between protocols, with iST and EasyPep showing highest efficiencies, followed by ISD-SDC protocols (**Supplemental Figure 2A**). The high efficiency of iST and EasyPep can be most likely explained by the combined use of trypsin and Lys-C in these kits, in contrast to trypsin alone as in the other protocols. This suggests that all methods could probably benefit from the use of both enzymes, which needs to be considered when comparing results across protocols.

The differences in cleavage efficiency also help to interpret the results of protein and peptide IDs (**Figure 2A**). Some methods with high peptide ID numbers show comparably lower protein IDs (e.g. U-A, FASP, SP3). However, when considering peptides with no missed cleavages (**Figure 2A**, lower panel) it is evident that the lower digestion efficiency in these methods might resulted in lower proteome coverage. The excellent performance of ISD-SDC protocols in terms of protein and peptide IDs even without additional use of Lys-C supports the originally reported properties of SDC to enhance trypsin activity and to increase digestion efficiency ^23^. Notably, the majority of cleanup protocols as well as the classical ISD urea protocols and SPEED showed good performance and rather similar numbers of IDs. Conversely, samples prepared in GnHCl-based buffers displayed the lowest numbers of protein and peptide IDs, suggesting interference of GnHCl with trypsin protease activity even at low concentrations, as has been reported before ^3,24^.

The values depicted in **Figure 2A** represent the sum of multiple effects, which hampers an independent evaluation of the impact of single method parameters, such as protein precipitation. To elucidate the unique impact of variables on the overall performance of ISD protocols, we applied linear regression modelling. In each model, the number of IDs were explained additively by the supposed independent effects of individual precipitation methods and buffer conditions, in addition to batch effects that derive from technical variance during the MS measurements (Supplemental Figure 2B). On the basis of model parameter estimates, we calculated protein and peptide IDs for individual precipitation strategies (Figures 2B) and buffer conditions (Figures 2C) that are corrected for the effects of all other model variables.

In general, protein precipitation only minimally affected the efficiency of protocols, with acetone and chloroform-methanol precipitation being slightly advantageous compared to the other methods (**Figure 2B**). The strongest impact on method performance is caused by the type of extraction buffer, which confirms that effective protein digestion is a key determinant for proteome coverage. It is possible that there are additional interaction effects between variables. For example, the bimodal data distribution in acetone precipitated samples could hint that acetone precipitation efficiency is influenced by buffer type. However, such potential effects are difficult to resolve statistically with the current study design and with the available number of datapoints and would require further and more specific experiments. Generally, the SDC-based buffer resulted in the highest numbers of identified proteins and peptides even without precipitation, whereas other methods like urea ISD clearly benefitted from precipitation protocols. Certainly, as mentioned before, these results have been obtained with HeLa cells and might not be directly translatable to other cells, tissues, or organisms with more challenging properties or specific requirements.

### Sample preparation artifacts

We next tested whether individual sample preparation methods are prone to protein modification-artifacts. We re-analyzed the MS raw data applying an open search strategy with the FragPipe proteomic software package ^25^. This software allows the assignment of delta-masses to peptides, which are indicative for modifications and adducts. We observed that the majority of PSMs (76 - 80 %) originated from unmodified peptides (**Figure 3A**). Most of the detected modifications were equally abundant in the different samples (**Figure 3B** and **Supplementary Table 2**), suggesting that they are either naturally occurring PTMs or inevitable, method-independent sample preparation artifacts. Nevertheless, we observed method-specific modifications and adducts, some of which have been previously described ^26–29^. Notably, all method-specific modifications were low in abundance (≤ 1%). For example, peptides in ISD-urea samples showed increased levels of carbamylation (**Figure 3C**), a well-known artifact for this buffer compound ^29^.

**Figure 3.**
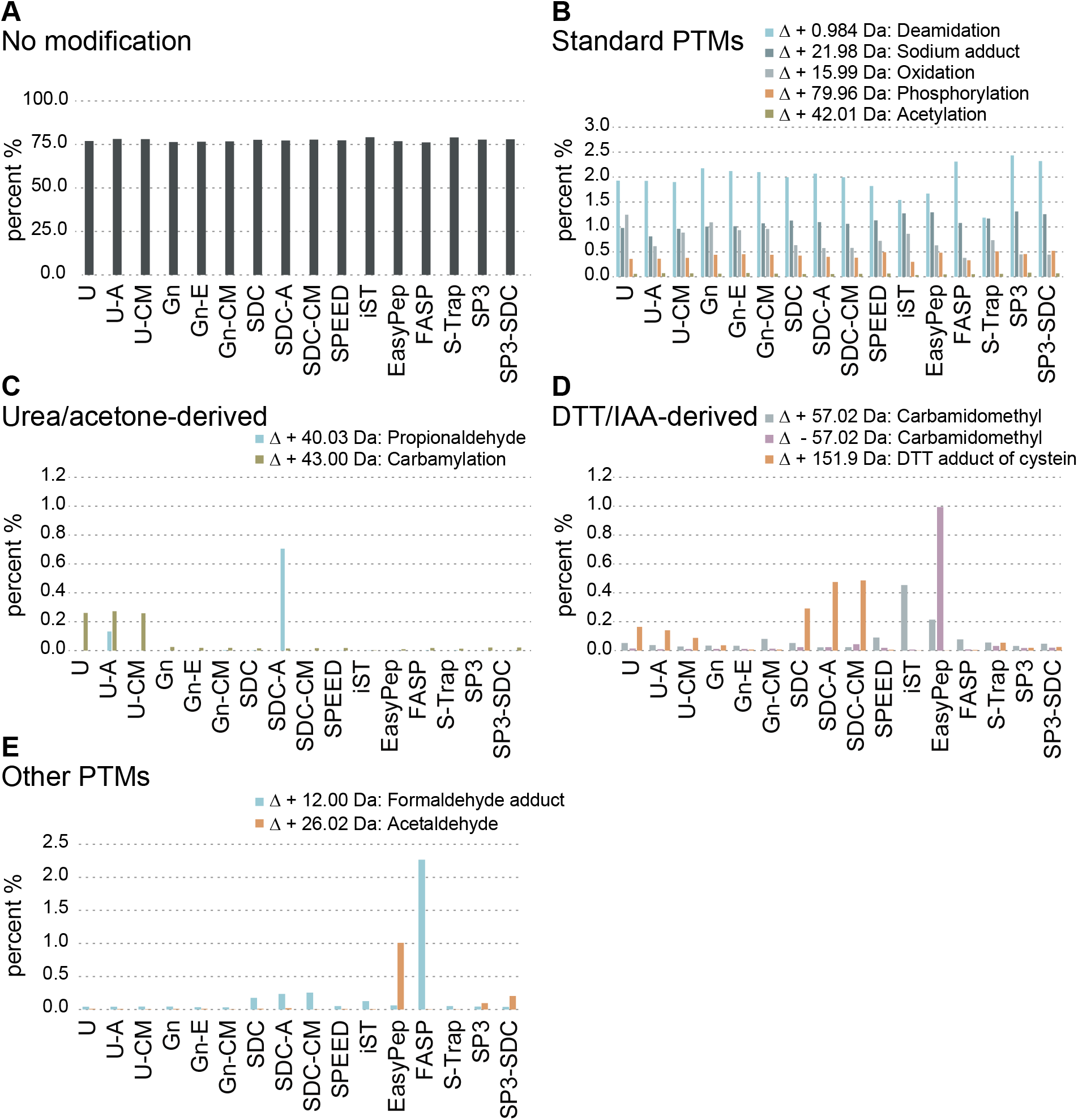
Analysis of covalent peptide modification artifacts created during sample preparation. Open search analysis using MSFragger to identify sample preparation-induced peptide modification artifacts. Barplots show the sum of PSMs of three replicates in percent (see also Supplemental Table 2). **(A)** Barplot showing percentage (y-axis) of peptides without modification. x-axis lists the applied sample preparation protocols. **(B)** Similar to (A) except that exemplary PTMs are shown. **(C)** Barplot highlighting previously described artifacts observed in samples prepared using urea-buffer (carbamylation) and acetone precipitation (delta mass: +40.03 Da), respectively. y-axis: percentage, x-axis: methods. **(D)** Similar to (C) except artifacts derived from reduction (DTT adduct of cysteine) and alkylation (carbamidomethyl) steps are shown. (**E**) Unknown modifications identified in EasyPep (delta mass: + 26.01 Da) and FASP (delta mass: + 12.00 Da). y-axis: percentage, x-axis: applied methods.

Peptide artifacts deriving from reduction and alkylation steps could be observed in several methods (**Figure 3D**). Despite reports on the disadvantages of using dithiothreitol (DTT) and iodoacetamide (IAA) we selected this protocol for the ISD as it is probably the most widely used and because it also allowed comparisons to other standard protocols such as FASP. Interestingly, the alkylation related artifacts were rather rare and appeared not as problematic as reported in literature ^30^. Although typical artifacts like off-target alkylation or DTT adducts could be detected, they were found to occur at low levels (< 0.5% or mostly lower), as also reported by Hains & Robsinson ^31^. Carbamidomethylated and carboxymethylated methionine or their according neutral losses ^30^ as well as potential dialkylation with IAA were not detected or occurred at levels below 0.01%. Among the minor effects, EasyPep and iST showed slightly elevated levels of off-target carbamidomethylation (+57.0215 Da predominantly on lysine and histidine) and EasyPep additionally for unmodified cysteines (−57.0215 Da), suggesting non optimal reaction conditions for alkylation of free thiols ^26^. Unfortunately, the type and concentration of chemicals used in these kits is not disclosed but based on the “one-pot” reaction conditions and published information ^13^ it can be assumed that chemicals other than IAA and DDT are used. Nevertheless, their impact on artifacts and general method performance appears to be rather small when compared to the other protocols in this study. ISD-SDC, and to a minor extent S-Trap, resulted in increased levels of DTT adducts on cysteine (+151.9966 Da). We further recorded a modification seemingly specific to acetone precipitation with a delta mass of +40.0313 Da (propionaldehyde) in ISD-urea and especially ISD-SDC samples (**Figure 3C**), possibly constituting acetone adducts ^28^. Finally, we observed enrichment of a delta mass +26.0157 Da in EasyPep samples likely corresponding to N-terminal acetaldehyde Schiff base formation ^27^ and a delta mass of +12.00 Da (formaldehyde adduct) previously described to be specific to FASP samples ^32^ (**Figure 3E**).

The open search strategy might not exhibit the sensitivity to reveal all modifications and artifacts occurring in the samples. However, it provides a rather unbiased, broad overview and revealed that only a negligible fraction of peptides was affected by method-induced modifications, indicating that artifacts induced by sample preparation pose only a minor problem for the protocols as they were applied in our study.

### Proteome coverage and qualitative differences

Apart from numbers of proteins and potential artifacts the most important question is certainly whether methods differ in terms of identity and quantity of the proteins they extract. We investigated whether the individual sample preparation methods covered largely similar or distinct fractions of the HeLa proteome (**Figure 4A**). Based on this analysis it appears that overall proteome coverage is rather comparable. We observed a predominant overlap of protein IDs when comparing classical ISD methods and SPEED (3498 proteins, 75.3% overlap). Similar observations were made when comparing the clean-up methods FASP, S-Trap and commercial kits EasyPep, iST (3711 proteins, 78.9% overlap) or when comparing SDC-A, FASP with SP3-based methods (3800 proteins, 81.9% overlap) (**Figure 4A**). The overlap of all 16 methods (2989 proteins) was 61.6% (**Supplemental Figure 3**) but this lack of overlap is certainly also driven to a large extent by missing identifications of rather low abundant peptides due to the stochastic nature of data-dependent acquisition. It is clear though, that a simple analysis of overlaps in protein IDs does not allow to reveal specific but more subtle differences.

**Figure 4.**
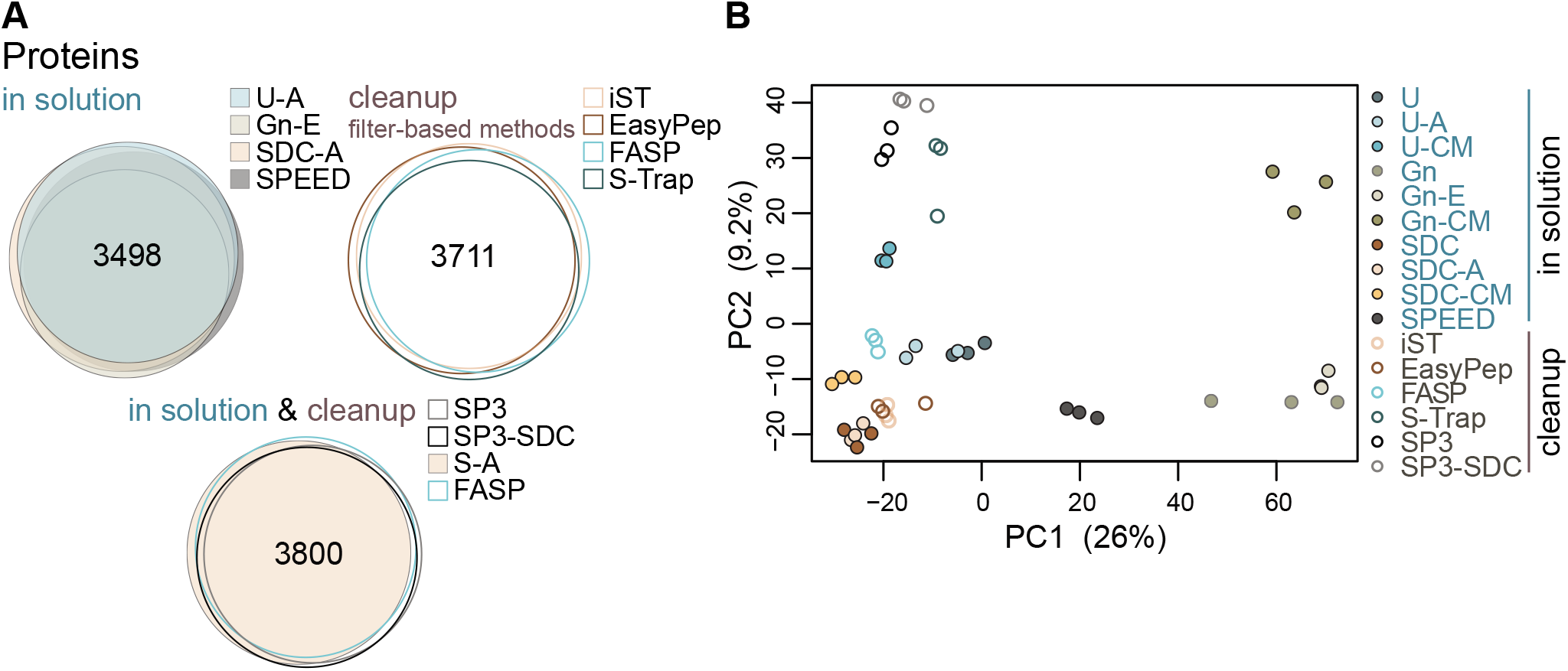
Overlaps of protein identifications and Principal Component Analysis. **(A)** Venn diagrams depicting the number of overlapping protein IDs (identified by MS/MS) obtained from different sample preparation methods. **(B)** Principal component analysis (PCA) based on log_2_ transformed LFQ intensities after normalization and imputation of missing values.

In contrast, a principal component analysis (PCA) of label-free quantified protein intensities separated out distinct clusters for the replicates corresponding to the different sample preparation methods, pointing towards qualitative differences in preparation-dependent variables (**Figure 4B**). We observed clustering according to buffer and precipitation conditions, with chloroform/methanol precipitation being more distant from other approaches. Distinct grouping of SP3-derived samples was also observed, irrespective of the applied buffer systems, suggesting that the magnetic bead-mediated protein pulldown poses a key variable for method-specific protein extraction. Furthermore, iST and EasyPep clustered close to SDC-ISD protocols, suggesting similarity in their methodology.

To further elucidate method-specific differences systematically, we carried out an explorative k-means cluster analysis and thereby classified variation patterns in protein intensities (**Figure 5A**). We first defined the optimal number of clusters, using the sum of squares within (SSW) distances to the next cluster center. Our approach defined nine k-means centers of cluster (k=9) as the optimal number, each showing a distinct method-dependent signature pattern of center-normalized LFQ intensities (**Supplemental Figure 4A**). Each cluster therefore consists of an individual set of protein IDs (**Figure 5B** and **Supplemental Figure 4B**). A downshift in center-normalized LFQ intensities suggests a method-dependent decrease in protein isolation efficacy in a given cluster. The opposite is true for observed upshifts. For clusters with a large number of elements, such as clusters 1 (n =1112) and 2 (n = 1935), we observed similar performance of all sample preparation methods (**Supplemental Figure 4B**). This suggests that the majority of proteins are effectively extracted independent of the applied protocol, which is also in agreement with the Venn diagrams (**Figure 4A**). Method-specific up- or downshifts in center-normalized LFQ intensities were prominent in clusters of smaller size, such as cluster 9 (n = 45) showing the most profound differences. Shifts in LFQ intensities were generally trending downward. **Figure 5C** summarizes the relative efficiency of sample preparation methods for each cluster in a heatmap (**Figure 5C**) and highlights that all methods display distinct profiles with unique features.

**Figure 5.**
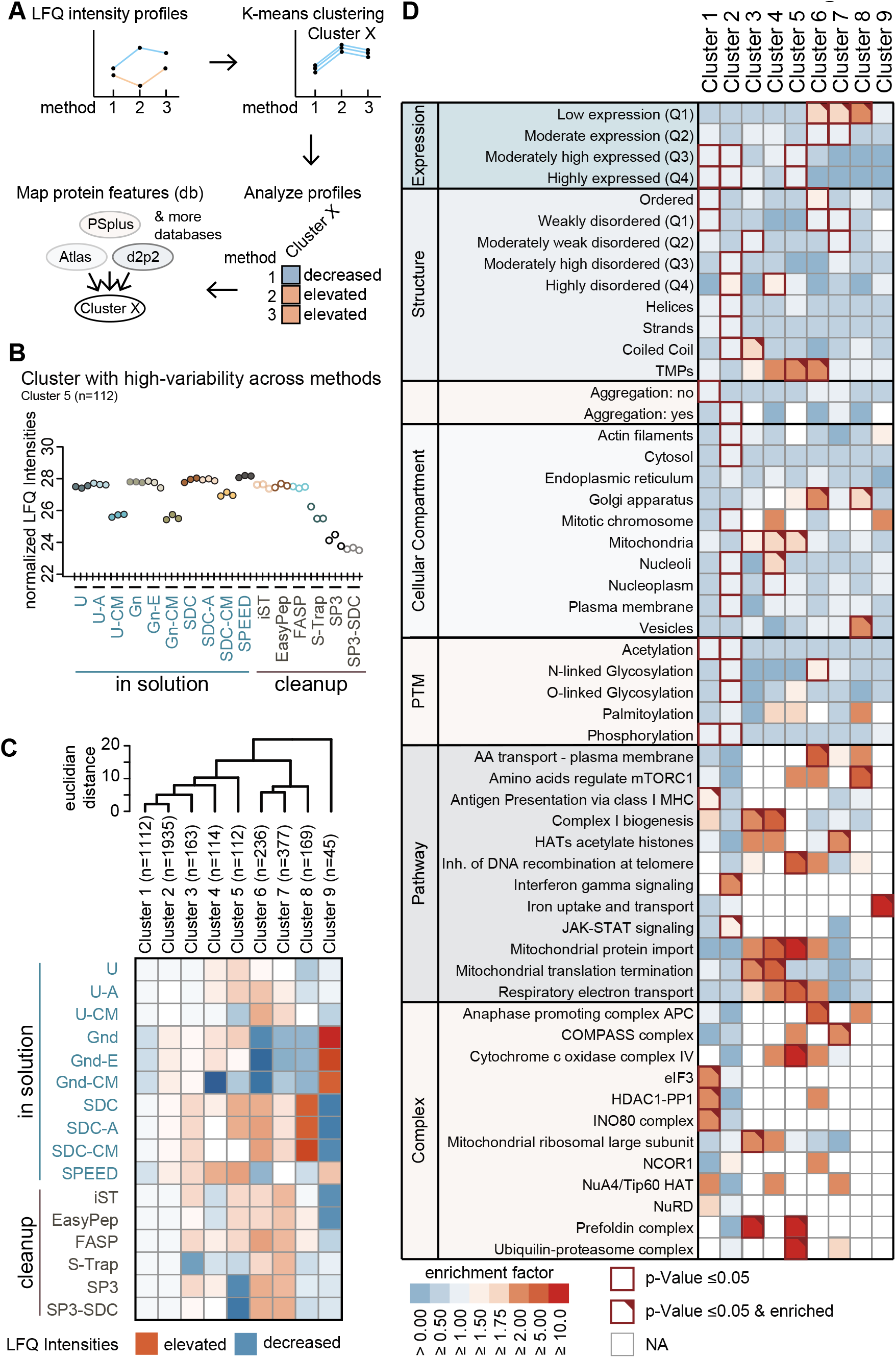
Exploratory k-Means cluster analysis and descriptive guide matrix. **(A)** Schematic illustration of k-means cluster analysis. **(B)** Representative cluster-specific (n = 112) profile plot of sample preparation methods resulting from an exploratory cluster analysis using k-Means. Methods (x-axis) are plotted against normalized log_2_-transformed LFQ intensities (y-axis). Plots depict the coordinates of k-means cluster centers. **(C)** Heatmap showing cluster specific deviations in the efficiency of sample preparation methods. Color-code represents average normalized log2 LFQ intensities. Dendrogram (top) depicts hierarchical relationships of clusters based on ultrametric euclidean distances. **(D)** Matrix depicting enrichment and significance of protein features (y-axis) in each k-means cluster (x-axis). Color code indicates enrichment factor of protein features. Red frames indicate significance (p-Value < 0.05). Red triangles: enrichment factor ≥ 2.

The clear clustering suggests that the respective proteins share common properties. We performed enrichment analysis on protein features that we extracted from selected databases, such as the Human Protein Atlas for subcellular localization ^33^, PhosphoSitePlus for known PTMs ^34^, PSIPRED for information on secondary structure ^35^, D2P2 providing a score for disordered regions ^36^, Pdbtm for transmembrane domains ^37,38^, Reactome.org for cellular pathways^39^, and three databases covering protein expression levels ^40^, complex information ^41^, and aggregation features ^42^ (**Figure 5A**). To determine which properties promote effective extraction by a given sample preparation method, we calculated cluster-specific enrichment for individual protein features (**Figure 5D** and **Supplemental Table 3**). As stated above, cluster 1 (n = 1112) comprises a high number of proteins that become efficiently isolated by all sample preparation methods (**Supplemental Figure 4B**). We found several features connected to the histone deacetylase (HDAC) 1 complex to be enriched in cluster 1, indicating the nuclear fraction of proteins can be purified with all tested methods at equal efficacy. Clusters 4 (n = 114) and 5 (n = 112) showed enrichment of several mitochondrion-associated properties, such as mitochondrial protein import, mitochondrial translation termination, respiratory electron transport, Cytochrome c oxidase complex IV, and mitochondrial ribosomal large subunit (**Figure 5D**). The fact that CM-based precipitation showed lower center-normalized LFQ intensity levels in clusters 4 and 5 (**Figure 5C** and **Supplemental Figure 4B**) suggests that these protocols should be avoided for mitochondrial proteomics. Conversely, ISD (without CM) and SPEED protocols seem to be well suited for mitochondrial protein extraction, as they resulted in the highest intensity levels (**Figures 5C, 5D** and **Supplemental Figure 4B**). Cluster 8 (n = 169) showed enrichment of vesicle- and membrane-associated protein properties (**Figure 5D**), which is consistent with the good performance of ISD-SDC in this group ^43^ (**Figure 5C**). Finally, proteins associated with iron uptake and transport were exclusively found to be enriched in cluster 9 (n = 45) (**Figure 5D**). Successful extraction of this set of proteins seems to be best achieved using ISD protocols based on GnHCl buffers.

Certainly, the efficacy of protein extraction of all applied methods could be further optimized. Here we provide a basis for doing so, indicating steps in sample preparation protocols that could be further fine-tuned. As suggested in previous reports ^1,4,11,17,18^ different combinations of buffer components and buffer systems, reactor types, proteolytic digestion protocols, and the use of nucleases could be implemented. Changes to protocols should, however, be made with caution, since cross-compatibility of reagents is not always guaranteed. For example, we occasionally observed gel-like phases in extracts when we used SDC in conjunction with phosphate-buffers (unpublished observation). Our data also suggests that omitting a protein precipitation step during MS sample preparation can still result in sufficient proteome coverage for HeLa cells. Yet, we generally advise to include a protein precipitation step to avoid carry-over of non-protein cellular components such as lipids, nucleic acids, metabolites etc. which could cause problems during later steps of sample preparation.

In general, different cell types and organisms may require different adaptations. To give an example, we observed that using buffer systems containing urea in combination with chloroform-methanol precipitation resulted in significant losses when proteins were extracted from *Saccharomyces cerevisiae* cells (unpublished observation). Doellinger *et al.*^4^ have shown that the SPEED protocol outperforms other protocols when processing bacterial samples. Furthermore, it is well known that samples from plants or fungi often require specific protocols due to the high-level of interfering metabolites.

Previous comparisons of sample preparation methods across species have shown that extraction bias does exist and that therefore a universal method is rather unlikely ^1,4^. Our study additionally demonstrates that even within the same sample type there is no one-fits-all protocol because all methods have their own peculiarities. For example, even though the SPEED protocol performs well in many aspects it also exhibits extraction bias towards certain groups of proteins (e.g. cluster 6). However, despite these clear differences for specific clusters our data also show that most methods, with the exception of GnHCl, perform overall rather similar in this cell type, which allows to choose methods rather on other parameters like ease-of-use, processing times, etc.

In summary, despite similar proteome coverage we could extract qualitative differences between the different protocols that represent varied purification efficacy for certain sets of proteins. The presented matrix, the underlying dataset, and the according methodology may serve as a guideline for the choice of a best-suited sample preparation method for a specific group of proteins of interest.

## Conclusion

The present study provides an in-depth and solid comparison of 16 of the most widely used MS sample preparation protocols in a human cell line. Careful attention has been paid to quality control and experimental design to maximize reproducibility, comparability, and to allow for unbiased statistical analyses. We demonstrate that the applied protocols had an overall rather similar performance with a low degree of protein modification artifacts and similar protein extraction efficiencies. Our analysis further revealed method specific protein clusters and we summarized their features in a guide matrix to assist in choosing an appropriate method. Urea-acetone, SDC-acetone and FASP protocols perform well in terms of number of covered protein/peptide IDs and enrichment of all classes of proteins. In addition, these methods are also comparatively cheap. A similar degree of performance was observed for the commercial kits, with the additional benefit that materials and reagents are provided in a standardized manner, and handling is straightforward. SPEED delivered in general a good performance and its simplicity and low price make it an attractive alternative but it seems this method could benefit from further refinements (e.g. trypsin & Lys-C digest). Other methods are preferable for enrichment for specific protein characteristics, such as ISD protocols based on GnHCl buffers for proteins associated with iron uptake and transport, however at the cost of reduced efficacy of proteolysis.

## Material & Methods

### Human cell culture

HeLa cells were cultivated in Dulbecco’s Modified Eagle’s Medium (DMEM 4.5 g/L glucose) (Sigma-Aldrich) supplemented with 10% fetal bovine serum (FCS) (Sigma-Aldrich), 1% L-glutamine (Sigma-Aldrich), 1% penicillin-streptomycin (Sigma-Aldrich) in 15 cm dishes under 5% CO_2_ at 37°C. Cells were harvested at ~ 80% confluency by 5 min treatment with trypsin (Sigma-Aldrich) at 37 °C, followed by a 1:1 dilution with full media to stop the digest. Cells were pelleted by centrifugation (5 min at 500 x g, 23°C) and washed with 1 x phosphate-buffered saline (PBS). Aliquots of 2.0E6 cells were subsequently snap-frozen in liquid N_2_ and kept at −80°C until lysis.

### In-solution protocols

#### In solution digests

HeLa cells (2.0E6 cells) were resolved in 100 μL denaturation buffer 0.1M Tris-HCl, pH 8.6, containing either 8M urea (U), 6M guanidine HCl (GnHCl), or 1% sodium deoxycholate (SDC), incubated for 10 min at room temperature (U) or at 60°C (GnHCl, SDC) in a ThermoMixer (Eppendorf), and subsequently disrupted by 2 × 20” high intensity sonication cycles at 4°C in a BioRuptor (Diagenode). Protein concentration was determined using the Micro BCA™ protein assay kit according to manufacturer’s instructions (Thermo Scientific). Each sample was split in two aliquots of 100 μg protein and one additional aliquot of 50 μg. Protein fraction of the two 100 μg aliquots were precipitated using acetone or chloroform-methanol, respectively. Only samples containing GnHCl were precipitated with ethanol instead of acetone, since GnHCl is not soluble in the latter. Protein pellets were dissolved in their respective denaturation buffer and protein concentration was determined as described above. Soluble proteins were reduced using 10 mM dithiothreitol (DTT) for 1 h at 37°C (U) or 60°C (SDC, GnHCl) and alkylated for 30 min using 20 mM iodoacetamide (IAA) in the dark. Chaotropic lysis buffers were then diluted to a final concentration of 1 M (urea) and 0.5 M (GnHCl). Proteins were digested overnight at 37°C, using trypsin (Trypsin Gold, Promega), in a 1:30 (w/w) enzyme to protein ratio. Digests were stopped by adding 10% trifluoroacetic acid (TFA) to a final concentration of 1%. SDC precipitates were removed by centrifugation (14,000 x g, 1 min, room temperature (RT)). 10 μg of resulting peptide samples were desalted on C18 StageTips (triple-plugs) ^44^, and eluted with 80% acetonitrile (ACN), 0.1% trifluoroacetic acid (TFA). After removal of elution buffer by vacuum centrifugation, samples were resuspended in 0.1% TFA, 2% ACN.

#### Sample Preparation by Easy Extraction and Digestion (SPEED)^4^

2.0E6 HeLa cells were resuspended in trifluoroacetic acid (TFA) (Merck) in a sample to TFA ratio of 1:4 (v/v), incubated at room temperature for 5 min, and neutralized with 2 M Tris Base using 8 x volume of TFA used for lysis. Reduction and alkylation of aliquots of 50 μg protein was achieved by incubation in 10 mM Tris(2-carboxyethyl)phosphine (TCEP) and 40 mM 2-Chloroacetamide (CAA) at 95°C for 5 min. Samples were diluted with ddH_2_O 1:5 and proteins were digested for 20 h at 37°C using trypsin (Trypsin Gold, Promega) at an enzyme/protein ratio of 1:50, as suggested in the original protocol. The digestion was stopped using 2% TFA (final concentration), peptides were desalted on C18 StageTips and eluted with 80% acetonitrile (ACN), 0.1% trifluoroacetic acid (TFA). Dried samples were resuspended in 0.1% TFA, 2% ACN.

### Device-based or cleanup protocols

#### Filter aided Sample Preparation (FASP)^8^

2.0E6 HeLa cells were resuspended in SDT-lysis buffer (4% SDS, 100 mM Tris-HCl, 100 mM DTT, pH 7.6) in a 1:10 (v/v) sample/buffer ratio, incubated at 95°C for 5 min and sonicated at 4°C for 2 cycles of 20 s at high intensity level using a BioRuptor (Diagenode). Samples were clarified by centrifugation at 16,000 x g for 15 min, at 24°C. Aliquots of 50 μg of protein were diluted in urea buffer UA (8M urea, 0.1 M Tris-HCl, pH 8.5) to a final concentration of 0.5% SDS. Protein extracts were further processed in a Microcon-30 kDa Centrifugal Filter Units (Merck) in a tempered centrifuge at 24°C. Samples were added to the filter unit, washed with UA buffer, centrifuged for 15 min at 14,000 x g, and incubated with 50 mM IAA for 20 min at room temperature (in the dark). SDS was exchanged by 4 consecutive washes with UA buffer (centrifugation: 15 min at 14,000 x g) and a single wash with 50 mM ammonium bicarbonate (ABC) followed by centrifugation for 5-10 min at 14,000 x g. Proteins were digested using trypsin (Trypsin Gold, Promega) in a 1:50 protein to enzyme ratio and incubation for 18h at 37°C on a thermo-shaker at 600 rpm. Resulting peptides were recovered by centrifugation at 14,000 x g for 5 min, followed by elution with 50 μL 50 mM ABC and repeated centrifugation. Combined eluates were acidified using TFA at a final concentration of 1%.

#### In-StageTip Sample Preparation (iST)

HeLa cell extracts were prepared using the iST 96x sample kit according to manufacturer’s instructions (PreOmics). In short, 2.0E6 cells were lysed by resuspension in lysis buffer solution at a target protein concentration of 1 mg/mL, heated to 95°C for 10 min shaking (1000 rpm) followed by 2 cycles à 20 s of sonication in BioRuptor (Diagenode). Aliquots containing 50 μg protein were transferred into a cartridge and cooled. The digestion solution was added and proteins were digested for 3 h at 37°C. Digestion was stopped by adding “Stop” solution and peptide purification was achieved by centrifugation for 3 min at 2250 x g, followed by 3 rounds of washing and elution into the collection plate using the provided solutions. Peptides were transferred to PCR tubes, dried in a vacuum-centrifuge, and resuspended in 0.1% TFA, 2% ACN for MS analysis.

#### EasyPep

HeLa cell extracts were prepared using the EasyPep™ Mini MS Sample Prep Kit (Thermo Fisher Scientific) according to manufacturer’s instructions. Briefly, 2.0E6 cells were lysed with Lysis Buffer aiming for a protein concentration of 1 mg/mL and aliquots containing 50 μg of protein were treated with Universal nuclease by 10 cycles of pipetting up and down until viscosity was reduced. Reduction and alkylation were achieved by addition of the respective solutions and incubation of samples at 95°C for 10 min. Once samples were cooled down, trypsin/LysC protease mixture was added and samples were digested for 3 h at 37°C. Tryptic digestion was stopped using “Digestion Stop Solution”. Peptide Cleanup columns were cleared from liquid by centrifugation and placed onto 2 mL microcentrifuge tubes. Sample mixtures were transferred into dry Peptide Cleanup columns. Two rounds of consecutive centrifugation and washing steps were performed. The columns were transferred to 2 mL microcentrifuge tubes and peptides were eluted by addition of Elution Solution and centrifugation at 1500 x g for 2 min. Samples were dried using a vacuum centrifuge and resuspended in 0.1% TFA, 2% ACN for MS analysis.

#### Suspension Trapping (S-Trap)^9^

2.0E6 HeLa cells were resuspended in lysis buffer LB (10% SDS (w/v), 0.1 M Tris-H3PO4, pH 7.55). Cells were disrupted by sonication (two cycles à 20 s at 4 °C) in a BioRuptor (Diagenode), and extracts were cleared by centrifugation at 15,000 x g for 1 min, 4°C. Aliquots of 50 μg protein were reduced by incubation with 20 mM (final concentration) DTT at 95 °C for 10 min and subsequently alkylated by addition of 40mM (final concentration) IAA and incubation for 30 min in the dark at room temperature. Samples were acidified with 1.2% (final concentration) phosphoric acid, mixed with 6 x volumes S-Trap binding buffer (90% MeOH in 0.1M Tris-H_3_PO_4_, pH 7.1), and loaded onto S-Trap columns that were placed in low binding tubes (Axygen). Solvent was removed by centrifugation (4,000 x g), proteins were washed three times with 150 μL S-Trap binding buffer and subsequently digested by addition of digestion buffer (500 mM ABC) containing 1:25 (w/w) trypsin (Trypsin Gold, Promega) and incubation at 37°C for three hours. Peptides were eluted in three consecutive steps by addition of 40 μl of 50 mM ABC, 40 μl 0.2% FA, and 35 μl 50% ACN, 0.2% FA followed by centrifugation at 4,000 x g, respectively. Eluates were pooled and concentrated in a SpeedVac (Thermo Fisher Scientific). Peptides were resolved in 0.1% TFA, 2% ACN. Aliquots of 10 μg of peptides were desalted on C18 StageTips (triple-plugs) ^44^, dried in a SpeedVac and resuspended in 0.1% TFA, 2% ACN.

#### Single-pot, solid-phase-enhanced sample preparation (SP3)^10^

Hela cells (2.0E6) were resolved in reconstitution buffer (RB) ^10^ or 1% SDC to a final protein concentration of 1 mg/mL and subsequently lysed, reduced (DTT 5 mM contained in RB) and alkylated using IAA (25 mM final concentration). For protein cleanup and digestion, samples of 50 μg protein were first mixed with SP3 beads in a 10:1 (w/w) beads-to-protein ratio. The mixture was then homogenized by adding 1 x volume of 100% EtOH and incubated for 5 min at 24°C shaking at 1000 rpm to induce protein binding to the beads. Proteins bound to beads were washed 4 x with 80% EtOH on a magnetic rack. On-bead digestion was achieved using trypsin (Trypsin Gold, Promega), added in a 1:30 (w/w) enzyme to protein ratio and 20 h incubation at 37□°C in a thermo shaker (shaking 1000 rpm). After digestion, beads were pelleted by centrifugation (20.000 x g, 1 min, 24°C) and supernatants containing peptides were transferred.

### Experimental design and quality control

To enable statistical analysis, we prepared three replicates of equal peptide concentration of each sample preparation method and applied several quality control steps which are summarized in detail below.

#### Type of replicates

Starting from a commonly cultured pool of HeLa cells three independent replicates were prepared for each sample preparation method. These replicates were defined as technical replicates.

#### Determination of protein concentration (of ISD samples)

Protein concentration after cell lysis and after protein precipitation was determined using the Micro BCA^™^ Protein assay kit (Thermo Scientific) according to manufacturer’s guidelines.

#### Determination of peptide concentration

An estimate of 250 ng of peptide per sample was mixed in 0.1% TFA, 2% ACN. Peptide concentrations were determined and adjusted according to UV chromatograms obtained at 214 nm on an UltiMate 3000 RSLC nano-HPLC System (Thermo Scientific), equipped with a monolithic column (PepSwift Monolithic RSLC, Thermo Scientific). To adjust peptide concentration for MS measurements, peaks were integrated using the chromatography software Chromeleon (Thermo Scientific) and peak areas were compared to in-house peptide standards of known concentration.

#### Equal loading of samples

All samples were adjusted to an estimated concentration of 100 ng/μL. Indexed retention time standard (iRT, Biognosys) was added to all samples to a final concentration of 0.1 injection equivalents (IE)/μL, allowing continuous monitoring of LC-MS/MS performance. Five μL of each sample corresponding to 500 ng peptide with 0.5 IE were subjected to MS analysis. Equal loading of samples was confirmed by checking total summed peptide intensities.

#### Organization of batches

Samples were organized into six batches. Batches 1-3 covered ISD protocols (including SPEED), with one replicate of each method per batch. Batches 4-6 were equally organized but with cleanup methods. Samples were measured in a randomized order, and all measurements were separated by wash runs. Before and after each batch, 25 ng of HeLa standard (Pierce) was injected to control system performance. Batches 1-3 and batches 4-6 were run in a single sequence, respectively.

#### (Post-acquisition) QC

Before and after each batch, 25 ng of HeLa standard (Pierce) were injected to control system performance. The quality of LC-MS runs was continuously monitored by checking the iRT signals in Skyline v20.1 ^45^. The number of missed cleavages and other metrics of quality control were determined using PTXQC ^46^.*Bridging of batches*: To account for changes in machine performance between batch sequences 1-3 and 4-6, three replicates of each group of batches (SDC-A and EasyPep, respectively) were re-measured in a single sequence of MS measurements. The differences in the number of IDs between these groups were ~1%, nevertheless the number of IDs of all original sample measurements were re-adjusted by the relative median change factor of the bridge samples. In short, the relative_median change_factor between the two groups was determined as [median (“SDC-A”_bridgesamples) - median (“EasyPep”_bridgesamples)] / median (“EasyPep”_bridgesamples). The corrected SDC-A median was calculated as [group_median (“EasyPep”_samples) + group_median (“EasyPep”_samples) * relative_median change_factor]. ISD groups were adjusted to the corrected SDC-A group median by their relative change to the original SDC-A group median.

### MS methods

LC-MS/MS analysis was performed on an UltiMate 3000 RSLC nano-HPLC System (Thermo Scientific), containing both a trapping column for peptide concentration (PepMap C18, 5 x 0.3 mm, 5 μm particle size) and an analytical column (PepMap C18, 500 x 0.075 mm, 2 μm particle size (Thermo Scientific), coupled to a Q Exactive HF-X Orbitrap (with HCD, higher-energy collisional dissociation mode) mass spectrometer via a Proxeon nanospray flex ion source (all Thermo Scientific). For peptide chromatography the concentration of organic solvent (acetonitrile) was increased linearly over 2 h from 1.6% to 28% in 0.1% formic acid at a flow rate of 230 nL/min. For acquisition of MS2 spectra the instrument was operated in data-dependent mode with dynamic exclusion enabled. The scan sequence began with an Orbitrap MS1 spectrum with the following parameters: resolution 120,000, scan range 375 – 1,500 m/z, automatic gain control (AGC) target 3□×□10^6^, and maximum injection time (IT) 60□ms. The top 20 precursors were selected for MS2 analysis (HCD) with the following parameters: resolution 15,000, AGC 1□×□10^5^, maximum IT 54□ms, isolation window 1.2 m/z, scan range 200 - 2000 m/z, and normalized collision energy (NCE) 28. The minimum AGC target was set at 1□×□10^4^, which corresponds to a 1.9□×□10^5^ intensity threshold. Peptide match was set to “preferred”. In addition, unassigned, singly and > 6+ charged species and isotopes were excluded from MS2 analysis and dynamic exclusion was set to 40 sec.

### MaxQuant Settings

Raw MS data was analyzed using MaxQuant ^47^ software version 1.6.14.0. MS2 spectra were searched against the canonical *Homo sapiens* (human) UniProt database (UP000005640_9606.fasta, release 2020_01, www.uniprot.com) containing 20607 entries, concatenated with the sequences of 397 common laboratory contaminants (extended MaxQuant contaminants database) and the iRTs. Enzyme specificity was set to “Trypsin/P”, the minimal peptide length was set to 7 and the maximum number of missed cleavages was set to 2. A maximum of 5 modifications per peptide was allowed. Carbamidomethylation of cysteine was searched as a fixed modification. “Acetyl (Protein N-term)” and “Oxidation (M)” were set as variable modifications. “Match between runs” and LFQ was activated. Results were filtered at a false discovery rate of 1% at protein and peptide spectrum match level.

### FragPipe analysis

Screening for protein modifications in an unbiased manner was performed using the open search option of MSFragger 3.3 in FragPipe (v16.0)^25^. All raw files were converted to mzML format using MSConvert ^48^ with peak picking activated. mzML files were assigned according to sample preparation methods and replicates in the Experiments/Group tab. Default open search parameters were used, with trypsin specificity, −150 to +500 Da precursor mass window, oxidation of methionine and protein N-terminal acetylation as variable modifications, and carbamidomethylation of cysteine as fixed modification. PTM-Shepherd was activated at default settings. The observed mass shifts were obtained from the “global.modsummary.tsv” and “global. profile. tsv” tables in the FragPipe output, inspected and filtered for abundant and relevant modifications.

### Computational methods

Computational analyses were performed using in-house R-scripts ^49^. The data was processed as follows: Proteins only identified by (modification) site, contaminant protein IDs as well as protein groups with less than two razor and unique peptides were removed and LFQ intensities were log_2_ transformed. Only IDs identified by MS/MS were considered. The data was filtered based on valid values in LFQ intensities with a cutoff of three valid values in at least one group. Remaining missing values were imputed by a constant equal to the minimal log_2_ LFQ intensity over all samples (rounded down). The data was normalized on average intensity. For principal component analysis (PCA) analysis the prcomp() function from the package stats (pre-installed in R) was used.

#### K-means clustering

K-means clustering was performed using the function kmeans() from the pre-installed R package stats. All the above described functions are embedded in the in-house script termed Cassiopeia ^49^. Briefly, Cassiopeia is an in-house built LaTeX script that runs on R-code and is used for the generation of quality control outputs, statistical outputs as well as for visualization of information for a given “proteinGroups.txt” file as produced by the quantitative proteomics software package MaxQuant ^47^.

#### Mapping of protein features

To map protein features, such as protein abundance levels, protein structure, localization in cellular compartments etc. to the clusters, results of the k-means cluster analysis have been merged with entries of protein databases using an in-house python script. The following databases have been used: Human Protein Atlas (proteinatlas.org ^33^), PhosphoSitePlus ^34^, PSIPRED^35^, D2P2^36^, Pdbtm ^37,38^, Reactome.org^39^, and a database covering protein expression level ^40^, information on complexes ^41^, and aggregator feature ^42^. Statistical significance for a protein characteristic’s enrichment in a cluster was inferred via one-sided Fisher’s exact test using the fisher.test() function from the R package stats. Enrichment factors were calculated as the ratio of observed number/expected number, where the expected number was calculated as the cluster size of cluster k multiplied by the relative frequency of the characteristic n throughout the whole experiment (i.e. enriched compared to global relative frequency).

#### Linear regression modelling

Within ISD-samples, the total number of observed features (proteins, peptides and peptides with 0 missed cleavages) were analyzed by means of a simple linear model applying the model formula: *IDs ~ batch + precipitation + buffer*. F-/ANOVA tests were applied to test for significance of the individual variables (batch, precipitation method and buffer). Linear model predictions were visualized as partial residual plots using the function effect_plot() from the R package jtools.

#### Venn diagrams and UpsetR-plots

MaxQuant ProteinGroups.txt output table was cleared from contaminants, reverse hits, ID’s only by site and iRT (internal retention time standards)-hits. Only protein IDs (protein groups) identified by MS/MS in at least two out of three replicates were considered. Area proportional Venn diagrams were created using the web application DeepVenn ^50^. Overlaps of protein ID’s (%) were quantified in Python (version 3.9) with pandas data analysis toolkit. UpSetR: Intersecting sets of protein IDs in all methods were further visualized in an UpSet plot, which was generated using a scalable matrix-based visualization script employed by the open source R package UpSetR ^51^.

### Cost-effort calculations

The financial expenditure for a method was determined by cost per sample either according to manufacturer (*e.g.,* 96 samples for iST 96x kit) or calculated by reagent cost per sample. For SP3, the cost was determined by the usage of magnetic beads solution per sample. The cost of one-time investments such as magnetic racks needed for SP3-protocols are not included in our table. The hands-on times refer to the processing time of 6 (12 samples for SP3 and SP3-SDC) samples in parallel without digestion, additional C18 clean-up or vacuum-centrifugation times.

The mass spectrometry proteomics data have been deposited to the ProteomeXchange Consortium via the PRIDE ^52^ partner repository with the dataset identifier PXD030406 and 10.6019/PXD030406.

## Acknowledgements

We thank Natalie Romanov for the help with the integration of protein databases used in the cluster analysis and for critical feedback on the manuscript, Dea Slade for kindly providing HeLa cells, Karl Mechtler and his team for the great collaborative spirit in our joint lab space, and the VBCF for providing the LC-MS instrument pool. GV was supported by FEMtech of the Austrian Forschungsförderungsgesellschaft (FFG). MM, NH, WR and MH were supported by the Austrian Science Fund (FWF) Special Research Program F70.

## Authors’ contribution

MH conceptualized the study. GV, DA, and MH designed experiments. GV, DA and NH performed experiments. GV, DA, MM, WR and MH analyzed the data. GV, WR, and MH wrote the paper. All authors edited the text. All authors read and approved the final manuscript.

## Supplemental Figure Legends

**Supplemental Figure 1 - Supplement to Figure 1. (A)** Scheme illustrating experimental workflow for ISD samples, including results from quality control testing. HeLa proteomes were extracted using urea (U)-, guanidine hydrochloride (G)-, or sodium deoxycholate (SDC)-buffered systems. Upper bar plot indicates extraction efficacy determined using the BCA™ Protein assay kit (Thermo Scientific). Lysates were either directly submitted to tryptic digestion or precipitated using acetone (A), ethanol (E) or chloroform/methanol (CM). Central bar plot: Precipitation efficacy was determined using the BCA™ Protein assay kit (Thermo Scientific). Tryptic digests were desalted using C18 stage tips and efficacy of proteolysis was determined by quantifications of UV chromatogram peak areas. Samples were adjusted to ensure equal loading for MS measurements. **(B)** Scheme illustrating experimental workflow for cleanup samples, including quantifications of UV chromatogram peak areas. **(C)** Box plots showing distributions of non-normalized log_2_ protein group intensities (top) as well as normalized log2 LFQ intensities (bottom) for each sample (y-axes). LC-MS batch numbers are indicated below samples (x-axis).

**Supplemental Figure 2 - Supplement to Figure 2.** (**A**) Bar diagram indicating number of single, double, triple missed cleavage peptides in percent. Diagram has been adapted from the PTXQC ^46^ report. (**B**) Partial Residual Plots depicting batch effects of MS measurements on the number of identified proteins (left), identified peptides (middle) and peptides with no missed cleavages (right). Data points represent the number of IDs. Error bars correspond to a 95% confidence interval (CI). Black lines indicate the average number of IDs.

**Supplemental Figure 3 - Supplement to Figure 4.** UpSet plot visualizing intersections of protein IDs extracted by individual sample preparation methods (sets) in a matrix layout. (Top) The x-axis shows intersections of set combinations through gray bars which are labeled with their respective intersection size. (Bottom) The overall set-size of a sample preparation method is listed on the bottom left. Next to each method (row), a black dot represents the inclusion of the respective set in an intersection (column). Sets that are not included in an intersection appear light gray.

**Supplemental Figure 4 - Supplement to Figure 5. (A)** Left: K-means plot: optimal number of clusters k was determined based on the total sum of squares within (SSW) for different k. Nine clusters were defined (s*ee material and methods*). Right: dendrogram of cluster centers as a result of an agglomerative clustering of k-means cluster centers with ultrametric euclidean distance. **(B)** Profiles of all k-means clusters. Dots represent method’s normalized LFQ intensities.

Supplemental Table Legends

**Supplemental Table 1 – Supplement to Figure 2.** Table listing total number of identified (by MS/MS) proteins (sheet 1), peptides (sheet 2) and peptides with no missed cleavages (sheet 3).

**Supplemental Table 2 - Supplement to Figure 3.** Results obtained from the open search with MSFragger analysis output table “global.modsummary.tsv”. Sum of PSMs of corresponding replicates of all samples are shown in numbers (left) and percent (right). Sheet 2: MSFragger output table “global.profile.tsv”.

**Supplemental Table 3 - Supplement to Figure 5.** Full matrix depicting enrichment and significance of protein features of the exploratory k-Means cluster analysis shown in Figure 5. Columns: k-means clusters 1 - 9. Rows: protein features in each k-means cluster. Color code indicates enrichment factor of protein features.

