## Supplementary figures and images for "In search of the universal method: a comparative survey of bottom-up proteomics sample preparation methods"

### Supplemental Figure 1

**Supplemental Figure 1**

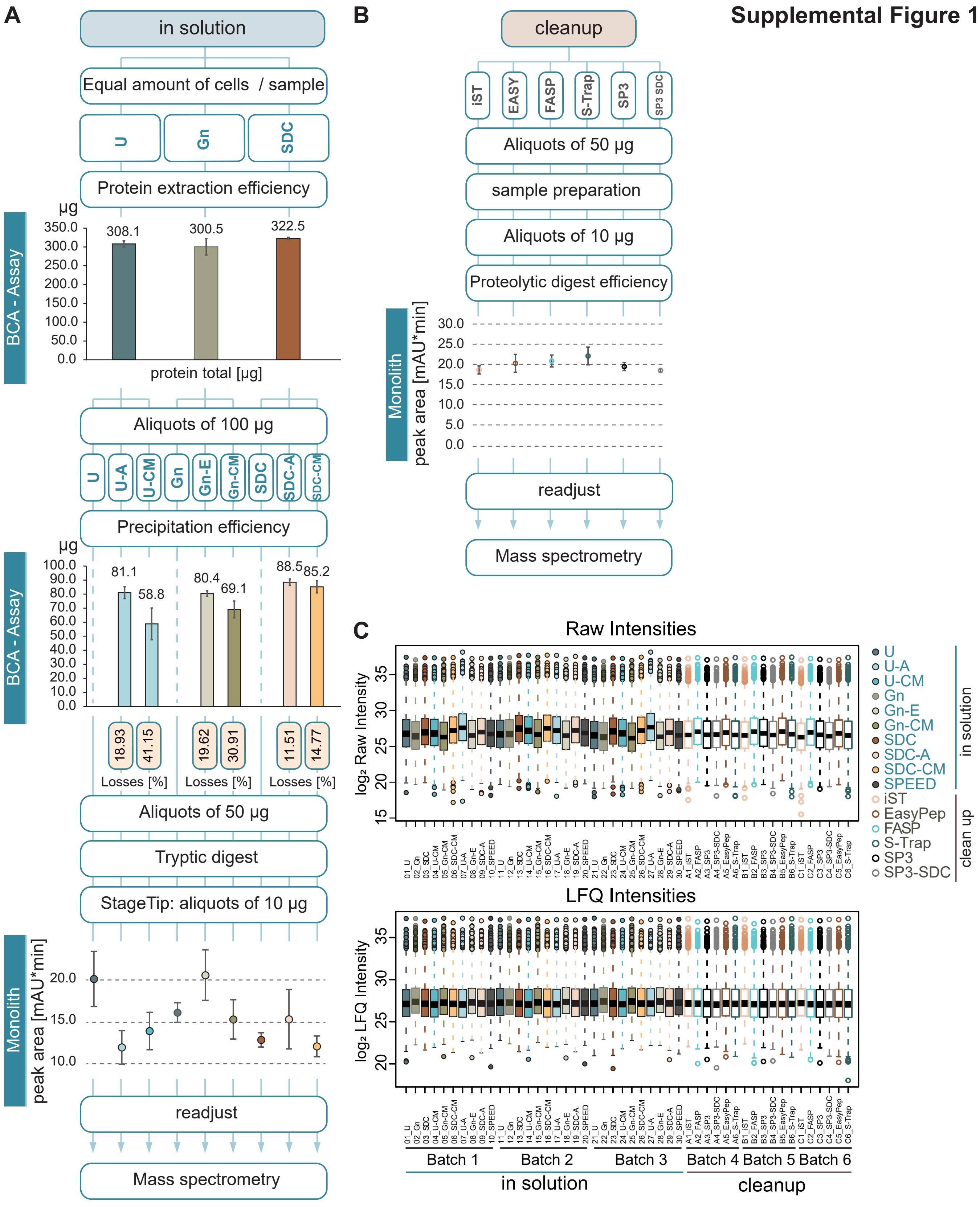

### Supplemental Figure 2

**A** **Supplemental Figure 2**

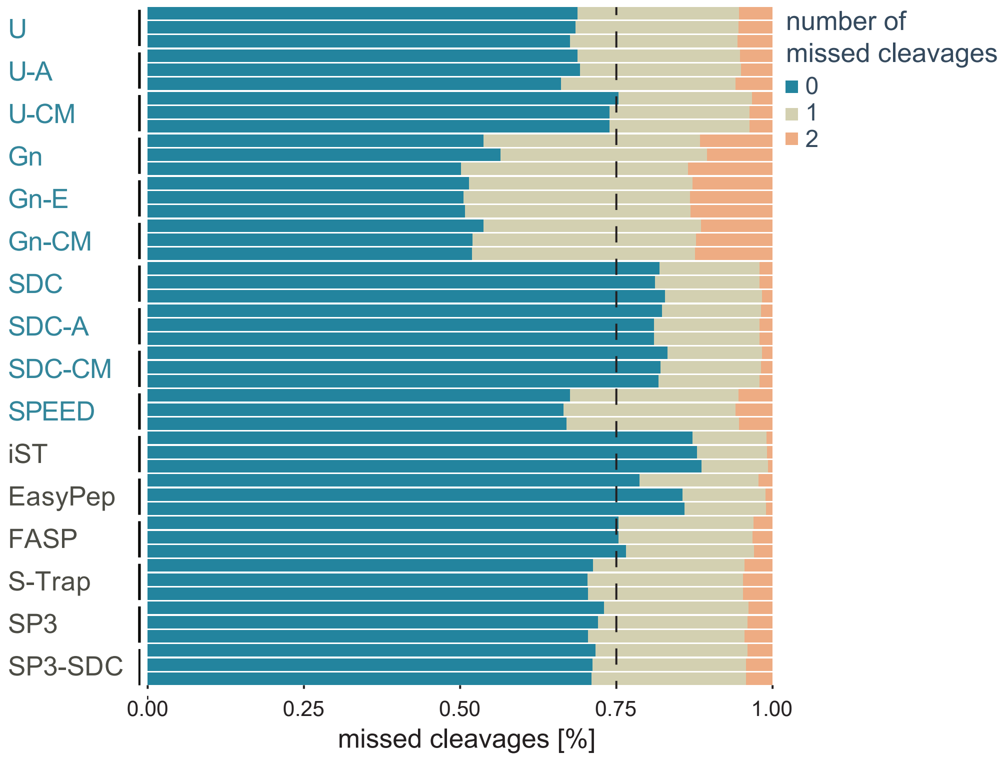


**B**

Batch effects - in solution

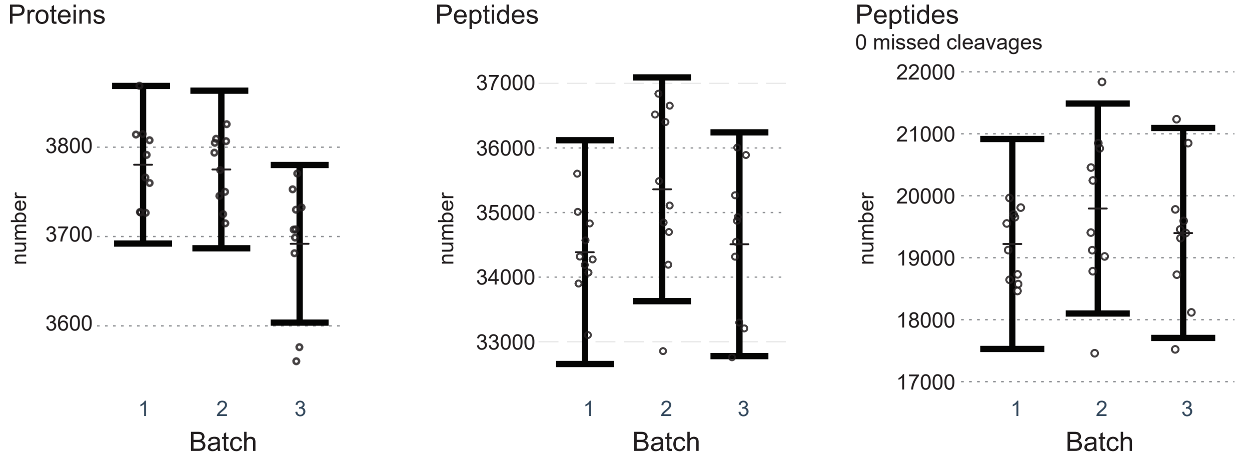

### Supplemental Figure 3

##### Supplemental Figure 3

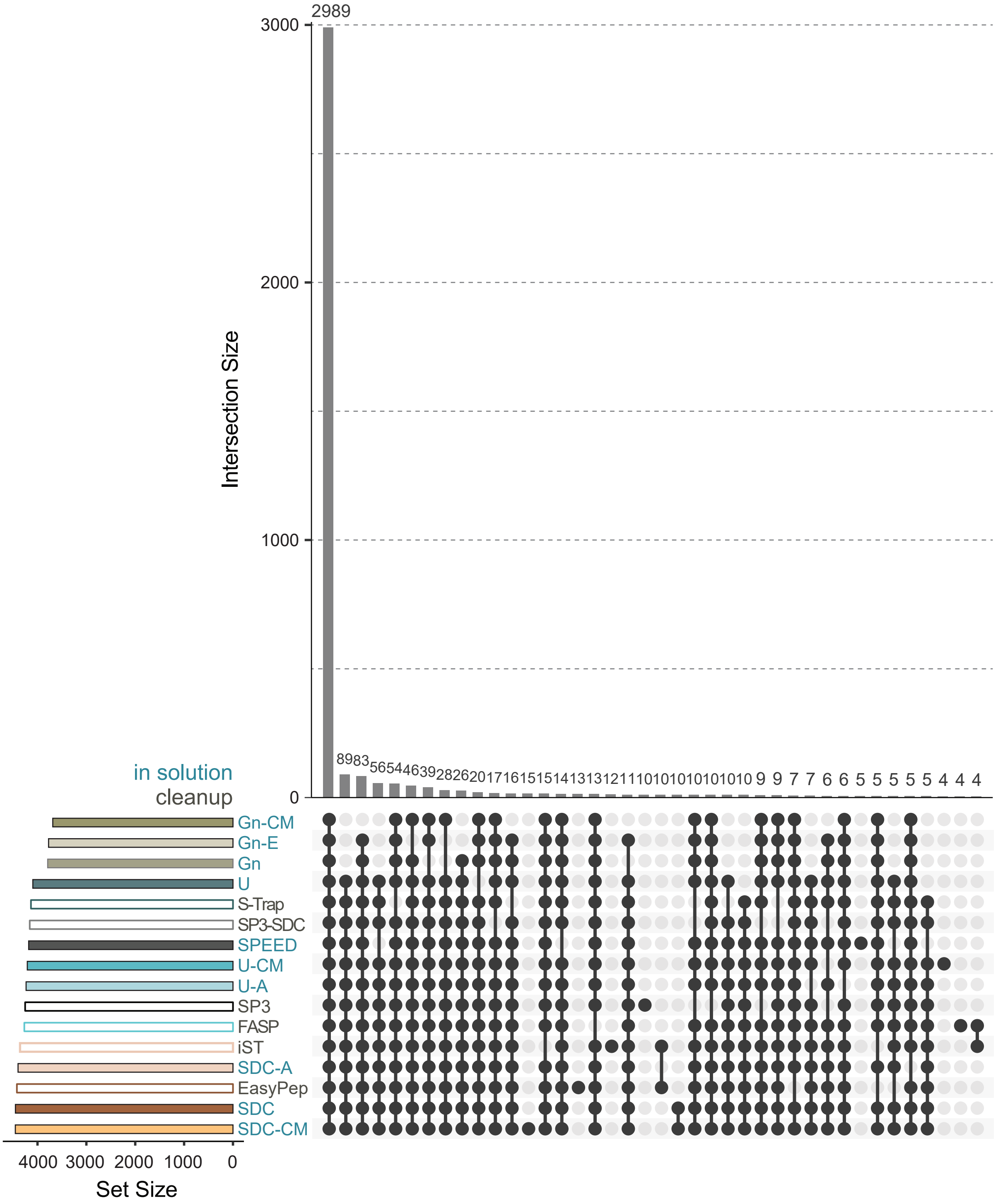

### Supplemental Figure 4

#### Supplemental Figure 4

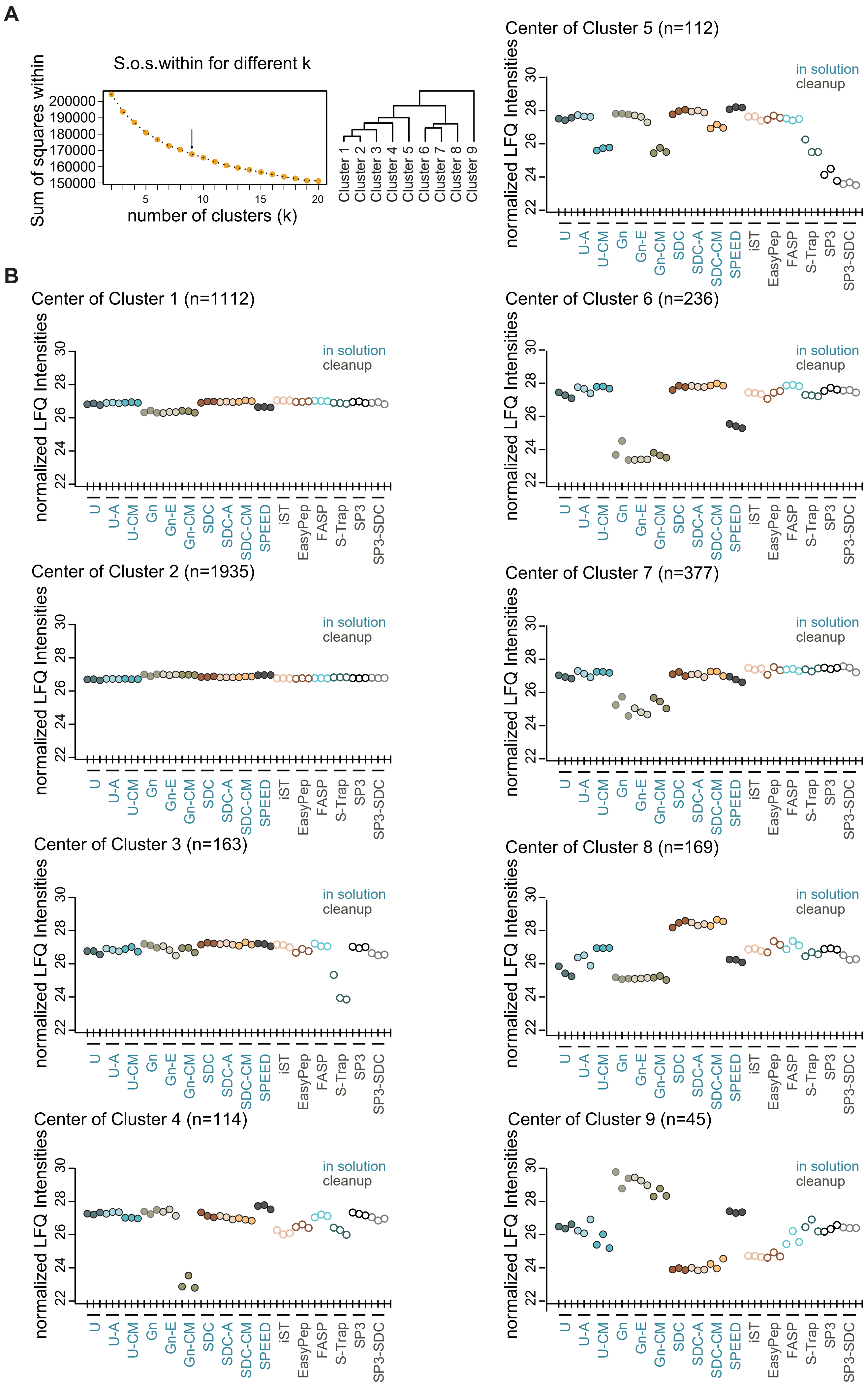
